# Beyond the Euclidean brain: inferring non-Euclidean latent trajectories from spike trains

**DOI:** 10.1101/2022.05.11.490308

**Authors:** Kristopher T. Jensen, David Liu, Ta-Chu Kao, Máté Lengyel, Guillaume Hennequin

## Abstract

Neuroscience faces a growing need for scalable data analysis methods that reduce the dimensionality of population recordings yet retain key aspects of the computation or behaviour. To extract interpretable latent trajectories from neural data, it is critical to embrace the inherent topology of the features of interest: head direction evolves on a ring or torus, 3D body rotations on the special orthogonal group, and navigation is best described in the intrinsic coordinates of the environment. Accordingly, we recently proposed the manifold Gaussian process latent variable model (mGPLVM) to simultaneously infer latent representations on non-Euclidean manifolds and how neurons are tuned to these representations. This probabilistic method generalizes previous Euclidean models and allows principled selection between candidate latent topologies. While powerful, mGPLVM makes two unjustified approximations that limit its practical applicability to neural datasets. First, consecutive latent states are assumed independent *a priori*, whereas behaviour is continuous in time. Second, its Gaussian noise model is inappropriate for positive integer spike counts. Previous work in Euclidean LVMs such as GPFA has shown significant improvements in performance when modeling such features appropriately (Jensen et al., 2021). Here, we extend mGPLVM by incorporating temporally continuous priors over latent states and flexible count-based noise models. This improves inference on synthetic data, avoiding negative spike count predictions and discontinuous jumps in latent trajectories. On real data, we also mitigate these pathologies while improving model fit compared to the original mGPLVM formulation. In summary, our extended mGPLVM provides a widely applicable tool for inferring (non-)Euclidean neural representations from large-scale, heterogeneous population recordings. We provide an efficient implementation in python, relying on recent advances in approximate inference to e.g. fit 10,000 time bins of recording for 100 neurons in five minutes on a single GPU.

## Manifold GPLVM

The details of mGPLVM are described by Jensen et al. (2020). Briefly, neural activity is assumed to arise from a set of latent states {*g*} on some potentially non-Euclidean manifold 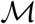 through a Gaussian process observation model defined on the manifold:

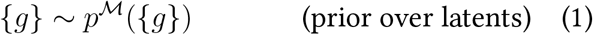

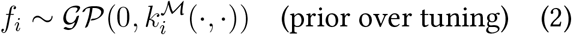

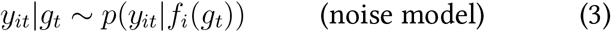

This model succesfully infers non-Euclidean latent states and tuning curves from synthetic and experimental data, and correctly identifies the underlying latent topology on manifolds ranging from rings and tori to the group of 3D rotations. Jensen et al. (2020) assume that the prior over latents (Equation 1) factorizes over time, 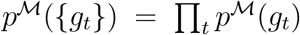 with 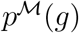 being uniform on 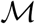. This is an inappropriate model for processes with known temporal dependencies such as navigation or recurrent network dynamics. However, extending the model to non-factorized priors is challenging due to the non-Euclidean nature of the latent space. Furthermore, the noise model in Equation 3 was previously assumed to be Gaussian, which is inappropriate for e.g. count-based data such as that arising from electrophysiological recordings. Here, we extend mGPLVM to both include temporally continuous priors and fit count-based observations.

### Continuous temporal priors

We start from the mGPLVM Evidence Lower Bound (ELBO; Jensen et al., 2020):

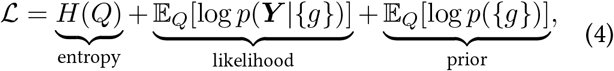

where *Q* is a variational distribution with parameters *θ*, and 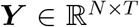 is the recorded activity of *N* neurons at *T* time steps. We encourage temporally continuous latent variable trajectories by introducing a Markovian prior:

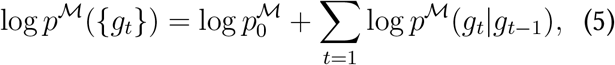

where the latent state at time *t* depends on that at *t* – 1. Here, we propose a tractable density 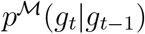 defined on Lie groups, which allows for Monte Carlo estimation of the prior. In Euclidean space, we can define a random walk prior as 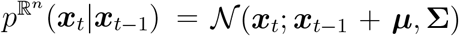, where ***μ*** and **Σ** are learnable parameters. Generalizing this to non-Euclidean Lie groups, we define a reference distribution on the tangent space, 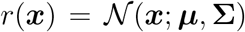, and project it onto the manifold:

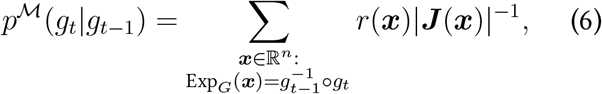

where ***J*** (***x***) is the Jacobian of the exponential map Exp*_G_* at ***x*** (Jensen et al., 2020; Falorsi et al., 2019), ***μ*** models any systematic drift on 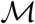, and **Σ** models the degree of continuity.

### Count-based observations

To generalize the observation model to fit count data, we use a variational distribution 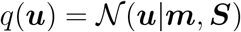 to lower-bound the GP likelihood term in the ELBO (Hensman et al., 2015):

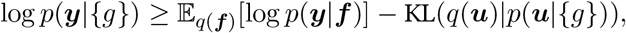

where ***y*** is a single row of ***Y, f*** are the corresponding function values of the GP, and

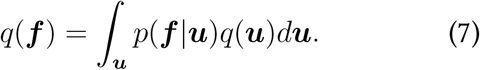

Under a Poisson noise model with an exponential link function commonly used for neural count data, this lower bound can be evaluated analytically. For more general noise models, we instead approximate 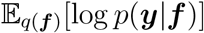 using Gauss-Hermite quadrature, which is applicable to any noise distribution with a closed-form likelihood *p*(***y***|***f***) (Hensman et al., 2015; Jensen et al., 2021). This allows us to fit more general count models and account for e.g under- or overdispersion in neural data with binomial or negative binomial noise models (Keeley et al., 2020; Liu and Lengyel, 2021).

## Results

For synthetic data with a smooth latent trajectory on a ring, mGPLVM recovers the true latents better with a time-continuous than a discontinuous prior (Figure A), and the learned parameters ***μ*** and **Σ** approximate the distribution over consecutive displacements. This leads to lower uncertainty in the variational distribution and improved log marginal likelihoods (ΔLL = (0.10 ± 0.02) × *NT*). For synthetic count data, the Poisson noise model improves the inferred tuning curves, capturing changes in uncertainty with preferred orientation and avoiding negative spike counts (Figure B). Combining these features, we compare models with temporally continuous or discontinuous priors and Gaussian or Poisson noise models on count data from a synthetic Poisson head direction circuit with trajectories generated from a random walk on the circle. We find that including both a continuous prior and Poisson noise improves the ability to infer ground truth latent states and reduces posterior uncertainty (Figure C). Our method also generalizes to higher dimensional non-Euclidean manifolds. We fit mGPLVM with a toroidal latent and time-continuous prior to count data from a 2D synthetic head direction circuit and recover ground truth latent states and tuning curves in a completely unsupervised manner (Figure D). Finally, we fit electrophysiological data from the mouse anterodorsal thalamic nucleus (Peyrache et al., 2015) using a circular latent. Neural firing is highly overdispersed for this data, and we find that a negative binomial noise model with a continuous prior outperforms a Poisson model and uncovers both the measured head direction and appropriate tuning curves (Figure E)

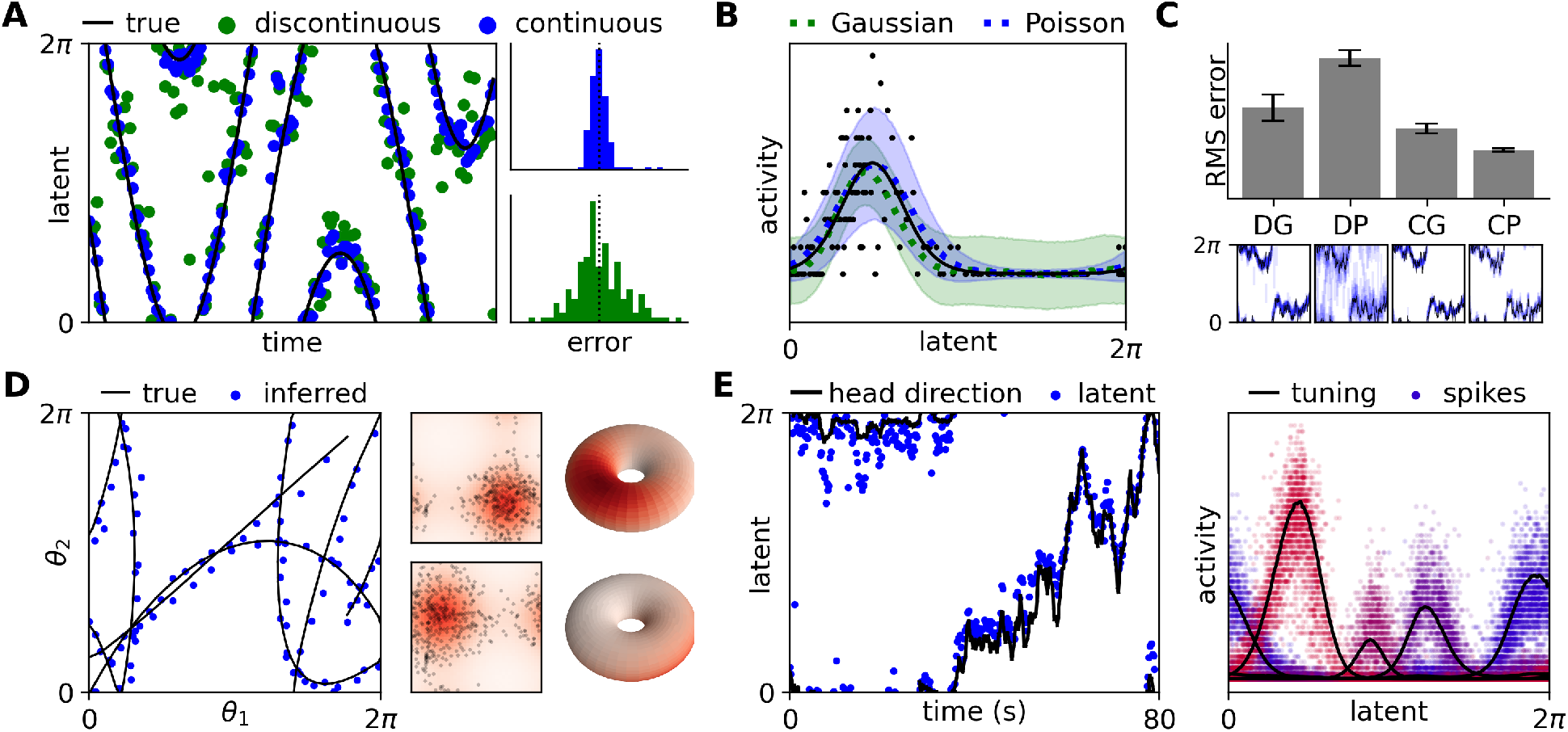
**(A)** True and inferred latents for mGPLVM fitted on a ring with time-discontinous (green) or -continuous (blue) priors. **(B)** Tuning curves for synthetic count data fitted with a Gaussian or Poisson noise model. **(C; top)** Recovery of latent trajectories with temporally discontinuous (D) or continuous (C) priors and Gaussian (G) or Poisson (P) noise models for synthetic head direction data (mean RMSE ± sem). **(C; bottom)** Example latent posteriors. Black lines are the ground truth and blue vertical lines indicate the posterior mean ± std. **(D)** Latent trajectory (left) and example tuning curves (right) for mGPLVM with a temporally continuous prior and Poisson noise fitted to synthetic count data on a torus. **(E)** Mouse head direction and inferred latents with a time-continuous prior and negative binomial noise model (left), and four example tuning curves (right).

